# Acute effects of drugs on *C.elegans* movement reveal complex responses and plasticity: Diversity of Acute Drug Responses in *C. elegans*

**DOI:** 10.1101/299719

**Authors:** Mark Spensley, Samantha Del Borrello, Djina Pajkic, Andrew G Fraser

**Affiliations:** Donnelly Centre for Cellular and Biomolecular Research, University of Toronto, Toronto, Ontario, Canada; Department of Molecular Genetics, University of Toronto, Toronto, Ontario, Canada

## Abstract

Many drugs act very rapidly — they can turn on or off their targets within minutes in a whole animal. What are the acute effects of drug treatment and how does an animal respond to these? We developed a simple assay to measure the acute effects of drugs on *C.elegans* movement and examined the effects of a range of compounds including neuroactive drugs, toxins, environmental stresses and novel compounds on worm movement over a time period of 3 hours. We find that many treatments show complex acute responses — a phase of rapid paralysis is followed by one or more recovery phases. The recoveries are not the result of some generic stress response but are specific to the drug e.g. recovery from paralysis due to a neuroactive drug requires neurotransmitter pathways whereas recovery from a metabolic inhibitor requires metabolic changes. Finally, we also find that acute responses can vary greatly across development and that there is extensive and complex natural variation in acute responses. In summary, acute responses are sensitive probes of the ability of biological networks to respond to drug treatment and these responses can reveal the action of unexplored pathways.

**Author Summary:** Drugs are powerful tools that let us switch on or off key pathways in whole animals and watch the effects. Here we set up a simple assay to measure how drugs affect the movement of the simple nematode *C.elegans* — crucially, we look how those responses change over time. We find that worms have complex responses to many different drugs — they go through clear phases of paralysis and recovery. The recovery from the initial effect of any drug is not due to a generic stress response but is specific to the way each drug acts. We find for example that worms can recover from paralysis driven by one neurotransmitter pathway by activating a different neurotransmitter pathway or from paralysis caused by loss of one metabolic pathway by turning on a second one. These complex responses show how the basic genetic networks that are needed for normal behaviour and function can rewire and respond to the effect of many drugs. Importantly, the responses can vary in many ways — different aged worms or different individuals can have different responses and capturing how drug responses change over time is critical to dissect this complexity.

## Introduction

Drugs are extremely powerful research tools. Addition of a drug can turn on or off a specific target protein and the effect on the organism can be followed over time. In model organisms, a well-characterized drug response can form the basis for genetic screens to identify the drug target and to find genes that modulate the effect of the drug. In *C.elegans*, for example, genetic screens for mutants with altered drug responses were key to finding the targets of several major anthelmintics (1–4) as well as to identifying core components of conserved neuronal signaling pathways (5,1,6,7).

Many genetic screens for *C.elegans* mutants with altered drug responses have screened for mutants that escape the effects of chronic exposure to drugs. In these screens, populations of mutant worms are typically exposed to drugs for several days (8–13). However, many drugs act very rapidly on worms yet currently relatively little is known about these acute responses of *C.elegans* to drugs. What do acute responses generally look like? Are acute responses the same at all developmental stages? Is there natural variation in acute responses as there is in mutant phenotypes or chronic drug responses? Do multiple drugs with the same long-term effect have the same acute responses or very different ones? What are the genes that affect the acute responses — are they the same or different genes as those that affect long-term drug effects?

Our goal in this study was to examine acute drug responses in *C.elegans* and address at least some of these questions. We developed an image-based method that accurately measures worm movement at high throughput. This allows us to gather rich information about the acute effects of drugs on worm movement, measuring both the effect of a range of doses of drugs and how the effects of any drug changes across time. We examined a wide range of drugs that were previously shown to be bioactive as well as a number of environmental stresses. We found that many drugs have complex acute effects on worms indicating that the animal responds rapidly to the initial drug effect — these acute responses include both transient and sustained recovery from initial paralysis. We show that the basis for complex acute responses is drug-specific and that acute responses can differ greatly between different developmental stages. Finally, we compare acute responses in two isolates and find that there is substantial variation in acute responses and that this changes with dose, time and developmental stage. We thus find that measuring acute responses to drugs can reveal the action of unexplored pathways and that acute responses vary extensively and at many levels between different individuals. The ability to measure acute responses at high throughput will allow both drug screens and genetic screens to uncover the underlying molecular basis for these complex responses.

## Results

### Development of a high throughput image-based movement assay

We wanted to measure the acute effects of drugs on worms over the time period of minutes to hours. We focused on movement as a key phenotype — a very wide range of compounds affect movement including toxins, neuroactive compounds and environmental stresses. There are a wide variety of assays to measure the acute effects of drugs on worm movement, with different levels of throughput and resolution, and different requirements for specialized equipment such as worm trackers or microfluidic devices (14–19). Here we focused on creating a simple assay for worm movement that allows us to follow how worms respond to drug treatment over time periods ranging from minutes to hours. The assay is simple, automated, gives quantitative data and has sufficient throughput to efficiently analyse responses to thousands of drugs.

In outline, our assay measures the movement of worms in 96-well format in liquid. Worms are continuously bathed in buffer containing the compound and there are multiple worms in each well (e.g. ~100 L1 larvae). We capture two successive images of each well, separated by a 500 millisecond interval — by comparing these two images we identify the extent of worm movement at any particular time point or drug concentration (see Fig 1A for details) and calculate a fractional mobility score (FMS) for each well at any individual time point. Worm movement can be measured at many time points to provide a time-course of the effect of a given drug concentration on worm movement (Fig 1B, right), at a range of drug concentrations to yield a dose-response curve (Fig 1B, lower) or (since the throughput is high enough) to cover both dose and time responses. The assay can thus be used to give a rich quantitative measurement of the effect of any drug on worm movement.

**Figure 1:**
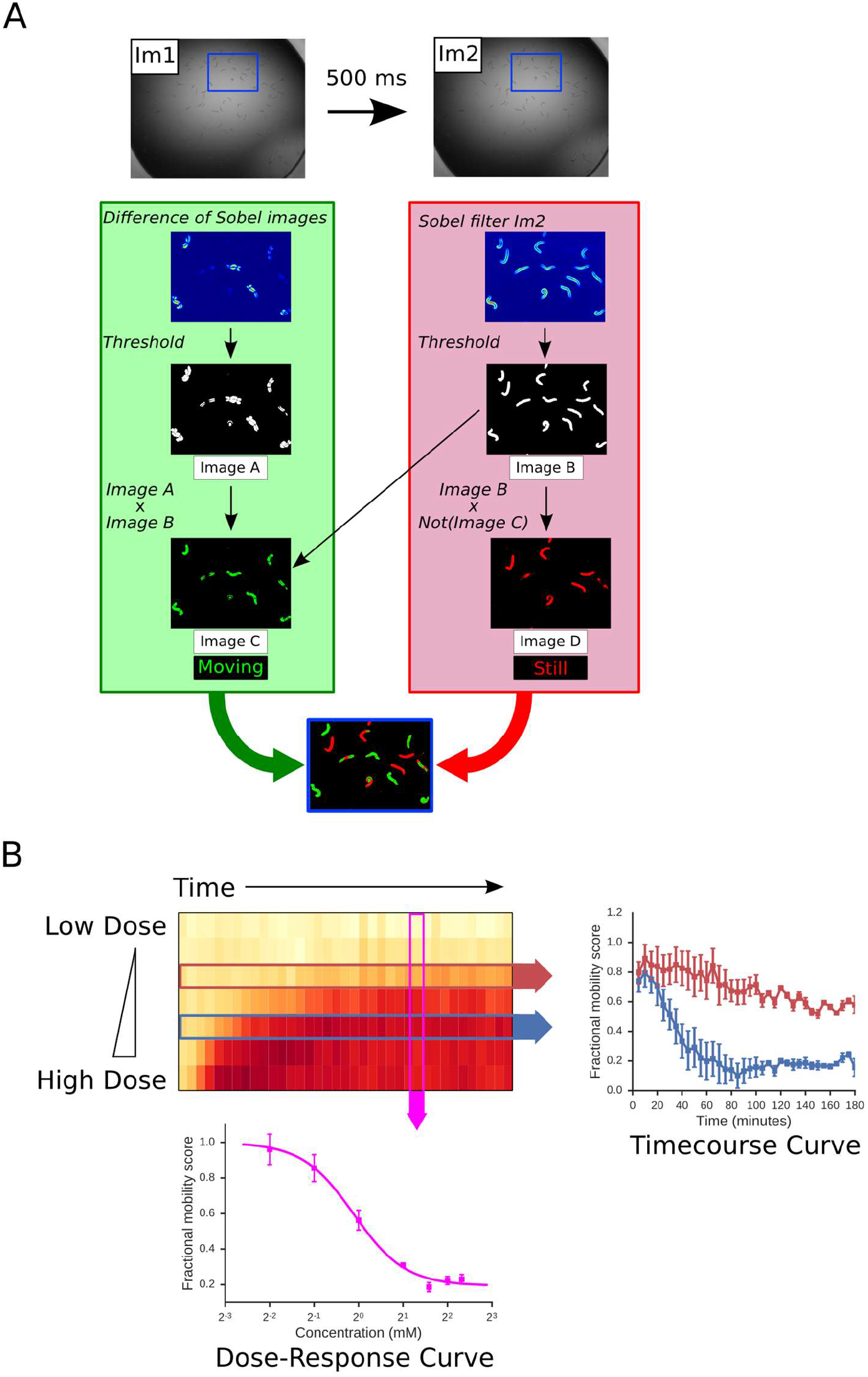
Outline of an image-based assay for worm mobility. **A:** Worms are placed in a 96 well plate in M9 buffer and treated with drugs as required. Two images of each well (Im1 and Im2) are captured at an interval of 500 ms at each timepoint. Images are processed as described in Materials and Methods to identify portions of the worms that move between the two images (green) and parts that remain stationary (red). This allows the calculation of a raw fractional mobility score (F.M.S.) as the fraction of total worm-associated pixels (red and green) that moved (green). This processing is illustrated in a magnified region of Im1 and Im2 (defined by blue box). **B: The image-based assay produces a rich view of drug effects on worm movement.** L4 worms were treated with a range of doses of aldicarb and movement measured at 5 minute time points over a 3 hour time course. The heat map shows the F.M.S. for each drug dose at each timepoint — white indicates full movement, red indicates lack of movement. Either dose response curves at a given time point (pink arrow) or kinetic responses to a single drug dose (red and blue arrows) can be extracted from these rich data. Data are means of four independent experiments. Error bars show standard error of the mean.

To validate our assay, and in particular to ensure that detected effects on movement are specific to the targeted pathway, we examined the response of adult worms to aldicarb (Ald), a very well characterised neuro-active compound (reviewed in depth in (20)). Acetylcholine (ACh) is a core neurotransmitter that is critical for normal movement (21). Following release of ACh at synapses, ACh is rapidly metabolised by acetylcholinesterases to turn off ACh responses. Aldicarb is a carbamate acetylcholinesterase inhibitor — Ald thus increases the local concentration of ACh at the synapse or neuromuscular junction by reducing the rate of degradation of endogenous released ACh (22). Aldi is known to cause paralysis in *C. elegans*, via the sustained activation of ionotropic acetylcholine receptors (nAChRs) and ACh signalling and the effects of both Ald and ACh have been extremely well studied (reviewed in (20)). We thus wanted to validate our assay by testing whether we can detect the well-established effects of Ald to induce paralysis and whether any Ald-induced paralysis is mediated by the well-characterised nAChR subunits, confirming that the effects we see are specific.

Consistent with previous studies, we found that Ald treatment results in paralysis of adult worms. Both the extent of paralysis and the rate of onset were dose dependent (Fig 2A). Aldicarb-induced paralysis is mediated by activation of nAChRs in body wall muscles; nAChRs are ligand gated ion channels composed of five subunits (23–26). Several of these subunits are known to be required for ACh-driven paralysis including UNC-38, UNC-63 and LEV-8. (23–25) To confirm that the effects of drugs on movement that we detect in our assay are specific to the activation of known AChRs (and not due to some kind of non-specific toxicity for example), we tested the response of mutant worms lacking functional copies of each of these subunits. As expected from previous studies, adult worms homozygous for loss-of-function mutations in *unc-38, unc-63, lev-8, lev-1* or *unc-29* are markedly less sensitive (Fig 2B, Fig S1). We also find that mecamylamine, a non-competitive nAChR antagonist, affects the dose-response relationship of aldicarb: in the presence of 600 μM mecamylamine, the EC50 of aldicarb shifts from 0.76 mM to 1.6 mM, with a slight decrease in maximal response (Fig 2C). The effects of Ald on worm movement that we measure in our assay are thus specific to the action of elevated ACh on nAChRs and not some non-specific effect on movement.

**Figure 2:**
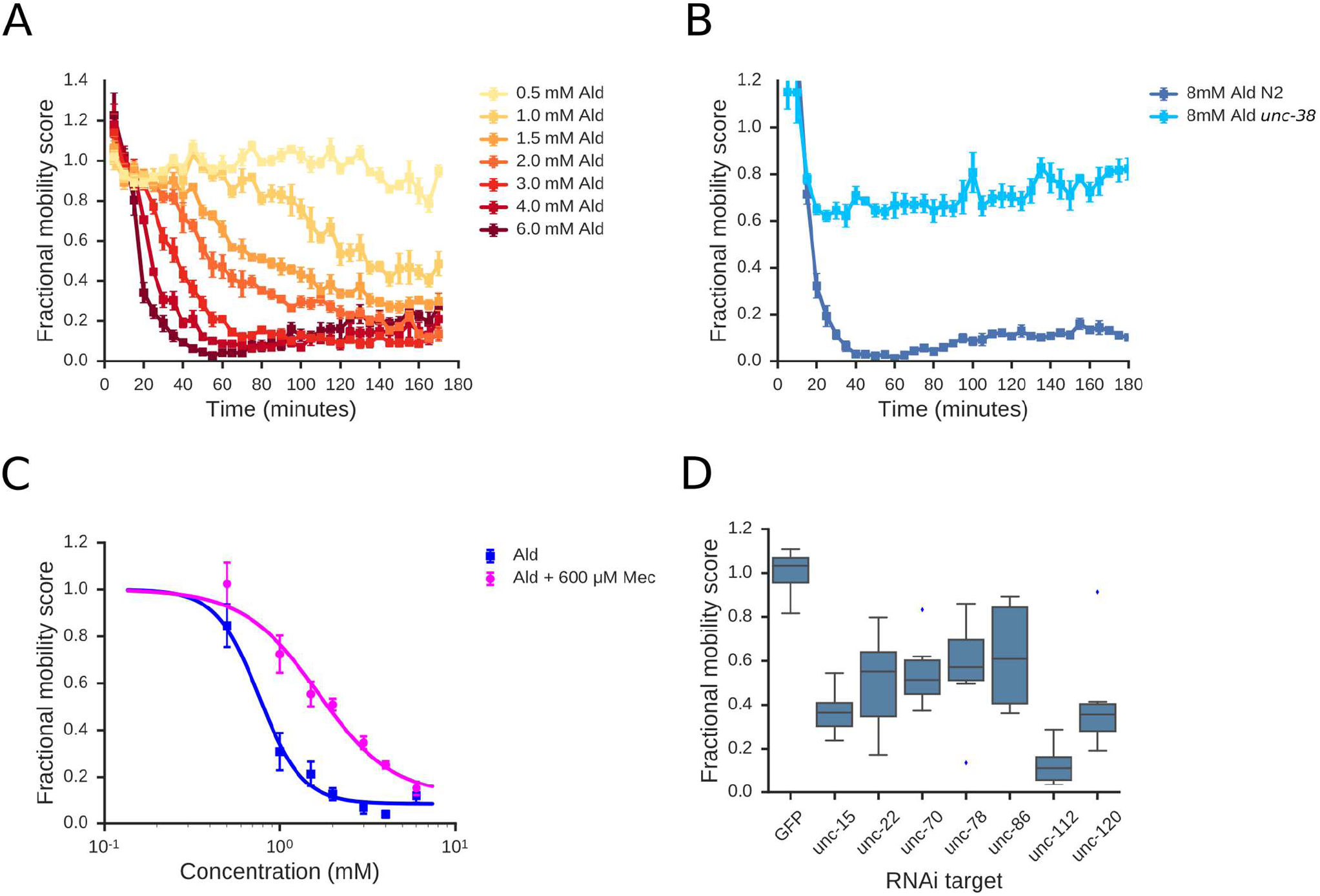
Assay characterisation. **A: Effect of aldicarb on adult worms**: Adult worms were treated with aldicarb at a range of concentrations and FMS was calculated at 5 minute intervals. FMS was plotted as a time-course of response for each concentration. **B: Adult worms homozygous for a loss-of-function *unc-38*(*e264*) mutation are insensitive to aldicarb.** A-C show means of 4 independent experiments; error bars show standard error of the mean. All data are normalised to drug-free controls.**C: The dose-response relationship for aldicarb is affected by addition of mecamylamine, an antagonist of nAChRs.** EC50 values were calculated by fitting a 3-parameter logistic function to the data (see Materials and Methods). **D: Assay detects effects of targeting genes required for normal movement using RNAi.** L1 worms were grown for 4 days in the presence of *E. coli* strains expressing dsRNA that target individual genes known to be required for normal movement. L1 progeny from these cultures were isolated by filtration and their mobility quantified using our assay. Mobility scores are normalized to a non-targeting RNAi control (dsRNA targeted against GFP). Data represent the mean of 8 wells, containing 100 worms per well.

We note that while we focus here on the use of this assay to measure the effects of drugs on worm movement, it can also be used to measure the effect of genetic perturbations on movement directly. To illustrate this, we used RNA-mediated interference (RNAi) to knock down 7 genes that are known to be required for wild-type movement and measured movement after RNAi. We found robust RNAi phenotypes for all 7 genes (Fig 2D) showing that our movement assay is applicable to both genetic and drug based screens. Thus, we expect our method to be applicable in a range of pharmacological, genetic, and RNAi experiments.

We have thus developed a simple method that allows us to measure the acute effects of drug treatments on worm movement at high throughput. We showed that our assay recapitulates the known effects of the well-characterised neuroactive drug Ald and that this is indeed due to its known mode of action through the activation of specific receptors. Crucially, our assay allows us to measure both dose responses and time-resolved responses to drugs. In the rest of this paper we demonstrate that time-resolved profiles of drug responses can yield new insights into *C. elegans* biology.

### Acute responses to drugs are often complex

We established a simple assay to measure acute responses of worms to drugs over time periods of minutes to hours. We wanted first to use this to explore acute responses in general — what do these look like for a variety of drugs and treatments? We exposed *C.elegans* L1 larvae to a wide range of different compounds and treatments including compounds that inhibit core essential machineries such as the electron transport chain (inhibition by potassium cyanide, KCN, shown in Fig 3A) or the ribosome (effect of cycloheximide in Fig 3B), environmental stresses such as high salt conditions (NaCl shown in Fig 3C), known nematicides (abamectin shown in Fig 3D), as well as compounds that affect movement via altered neurotransmission (e.g. aldicarb Fig 2A). In each case, worms were treated and their movement examined over a time course of 3 hours — at least 3 biological replicates were done for each treatment.

**Figure 3:**
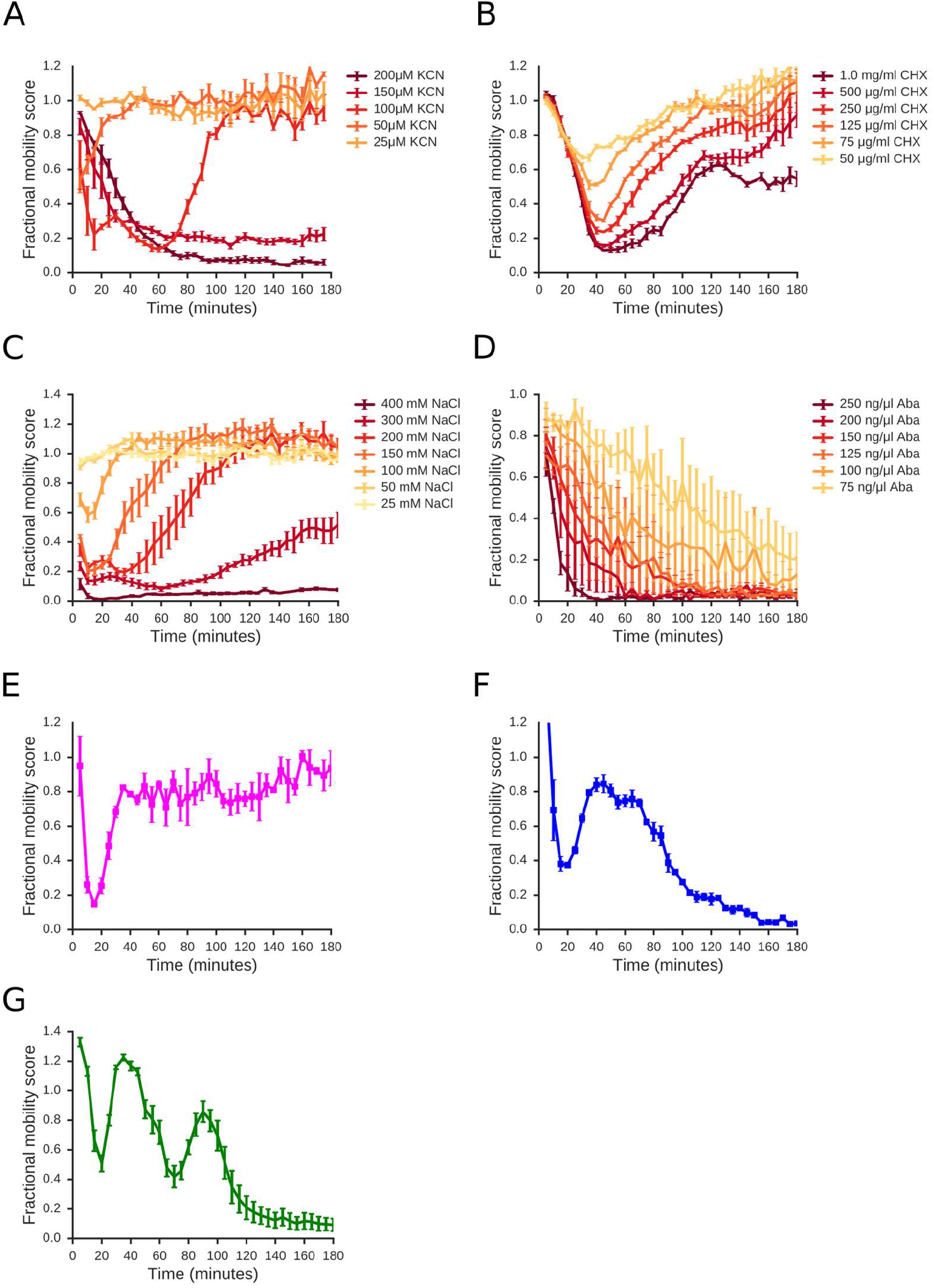
A wide range of compounds elicit movement responses. L1 worms were exposed to a range of doses of various compounds and movement quantified. **A: Effect of potassium cyanide on L1 worms.** Data are means of 4 independent replicates. **B: Effect of cycloheximide on L1 worms.** Data are means of 3 independent replicates. **C: Effect of sodium chloride on L1 worms.** Data are means of 4 independent replicates. **D: Effect of abamectin on L1 worms.** Data are means of 4 independent replicates. **E-G: Novel compounds elicit a range of dynamic responses.** Time-resolved responses for a number of novel compounds elicited a range of different response dynamics. L1 worms were continuously exposed to 40 μM of each compound. Data are means of 3 independent replicates. **(A-G)** Error bars show standard error of the mean.

In addition to this panel of characterized compounds, we also tested a range of novel compounds that had been identified as affecting worm growth in chronic exposure, population growth assays (13). We examined the effects of 170 of these on the movement of L1 larvae and find that many (53/170) also have acute effects on worm movement. Crucially, an unexpectedly high proportion (31 of 53 with acute effects; 58%) of drugs tested show complex time-resolved responses illustrated in Fig 3E-G, including either sustained or transient recovery from initial drug effects.

Having looked at acute responses to across a range of different treatments, we found complex acute responses across every type of compound, including neuro-active compounds, environmental stresses, and toxins that target essential genes. These responses can be very complex — for example, the novel compound shown in Fig 3G has two distinct phases of recovery and several structurally related compounds have similarly complex responses (data not shown). We note that every one of these complex drug responses can be the basis either for a genetic screen for mutants with altered responses or for drug screens for compounds that modulate the response. This complexity of many acute drug responses underlines the need to be able to measure time-resolved acute effects of drugs on movement.

The complex effects of many compounds over time suggest that the worms are responding to many drug treatments in some way — for example, several drugs cause an initial rapid paralysis followed by a recovery phase where the worms begin to move again at almost normal rates. What underlies this recovery? Is it some generic stress or xenobiotic response or does the worm respond in specific ways to each compound? To gain some insights into this we focused on the acute responses to two well-characterized compounds, Aldicarb and cyanide.

### Agonists of acetylcholine signalling have complex effects on worm movement that change across development

As shown in Fig 2A treatment of adult animals with Ald results in a rapid reduction in movement and sustained paralysis. Interestingly, we found that L1 animals behave completely differently— Ald induces rapid paralysis of L1 worms but this is followed by a more gradual recovery (Fig 4A; S2). To further examine the Ald response across development, we tested all developmental stages and found the response is qualitatively very different across development (Fig 4A). L1 and L2 stage larvae show an initial phase where Ald treatment results in reduced movement, followed by a recovery phase during which paralysis is gradually relieved. As worms progress through the four larval stages, the initial paralysis remains similar but this recovery is reduced, becoming minimal by adulthood. What underlies this recovery in L1 and L2 animals?

**Figure 4:**
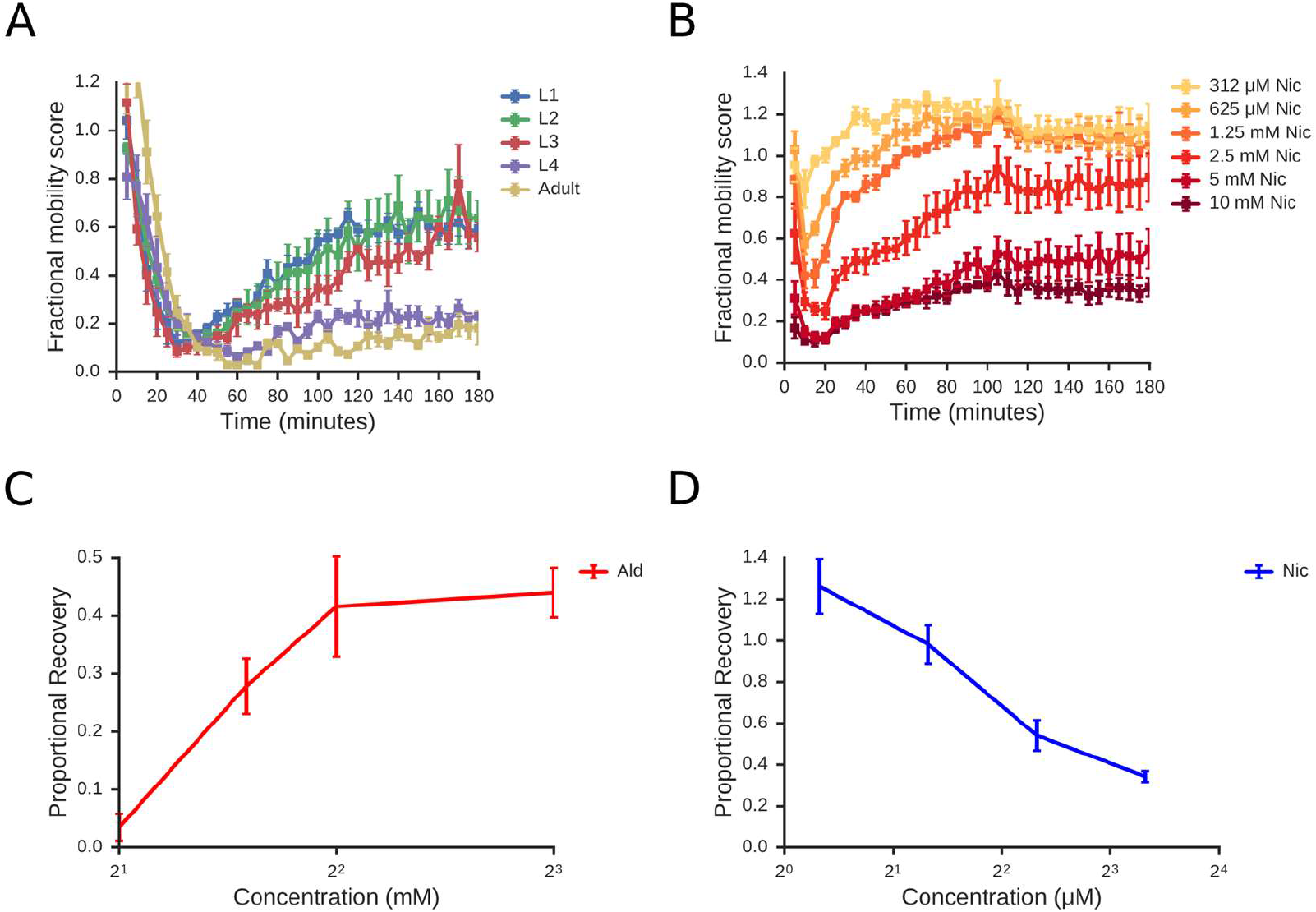
Larval worms show dose-dependent recovery from cholinergic paralysis. **A: Timecourse of response to 4mM aldicarb at different developmental stages.** Worms from each developmental stage were purified and dispensed using a COPAS biosort worm sorter before exposure to aldicarb as in Figure 2. Early larval stages show a robust recovery from paralysis, whereas L4 and adult worms show little recovery. Data are means of 3 independent experiments. **B: Response of L1 worms to nicotine.** L1 worms were treated with a range of nicotine concentrations and FMS measured over a 3hr time course. Data are means of 4 independent experiments. **C: Recovery of L1 worms from Ald-induced paralysis increases with higher doses.** For each concentration shown, a proportional recovery value was calculated as described in Material and Methods. Data are means of four independent experiments. **D: Recovery of L1 worms from Nic-induced paralysis decrease with higher doses.** Proportional recovery scores were calculated as above. Data are means of three independent experiments. **A-D:** Error bars show standard error of the mean.

To further explor the recovery of L1s from paralysis, we asked whether this is specific to Ald or whether other drugs that affect ACh signaling show similar acute responses. Ald affects ACh signalling by preventing the degradation of ACh — it does not activate ACh receptors directly nor does its action have any specificity for any specific sub-type of ACh receptor. Nicotine (Nic) activates ACh signaling in a different way — it is a direct agonist of specific ACh receptors, the nicotinic ligand-gated ion channels. We compared their acute responses and find that both Ald and Nic show a rapid initial paralysis followed by a recovery of movement (Fig 4B). While these two responses have superficial similarities, there is a crucial difference in the recovery phase. For Nic, while L1 worms recover from the reduction of movement caused by low doses of Nic, increasing concentrations of Nic result in greater paralysis and less recovery until at high Nic doses there is no appreciable recovery (Fig 4D). For Ald, we see the precise opposite — there is no recovery at low doses of Ald, but recovery increases with increasing Ald (Fig 4C). This suggests that the recovery from Ald and Nic-induced paralysis is fundamentally different and that Ald is somehow driving recovery since the more Ald we add, the stronger the recovery. How could Ald be acting and how could this be different to Nic?

ACh is known to activate two completely different classes of receptor: ligand-gated ion channels (nAChRs) and muscarinic ACh receptors (mAChRs). nAChRs act rapidly and are stimulated directly by Nic and indirectly by Ald via its action to increase ACh. mAChRs have much more complex G-protein coupled signaling — crucially, they are not activated by Nic but are also activated following Ald treatment. We hypothesized that the Ald recovery phase might be mediated by mAChRs and test this below.

### Recovery from rapid paralysis induced by Aldicarb occurs via muscarinic receptor signaling

To test whether recovery from Ald-induced paralysis might be due to the activation of mAChRs, we examined whether a mAChR antagonist, atropine (Atr), could block the recovery phase of the Ald response. This is indeed the case (Fig 5A) suggesting that recovery is the result of an activation of mAChR signaling following increased ACh levels after Ald treatment. The *C. elegans* genome encodes three muscarinic acetylcholine receptors, GAR-1, GAR-2 and GAR-3 (27–29) — we obtained deletion mutants for each of these receptors and tested whether the effect Ald was altered in these mutant strains. The Ald response is clearly different in the *gar-3(gk305*) mutant strain: while the paralysis phase appears very similar to wild-type, the recovery phase is strongly suppressed in the *gar-3(gk305*) mutant, although not completely abolished (Fig 5B). We note that mutations in *gar-1* and *gar-2* also have effects on recovery but these are much weaker and we suggest that *gar-3* is the key mAChR that drives recovery from nAChR stimulation in L1 animals.

**Figure 5:**
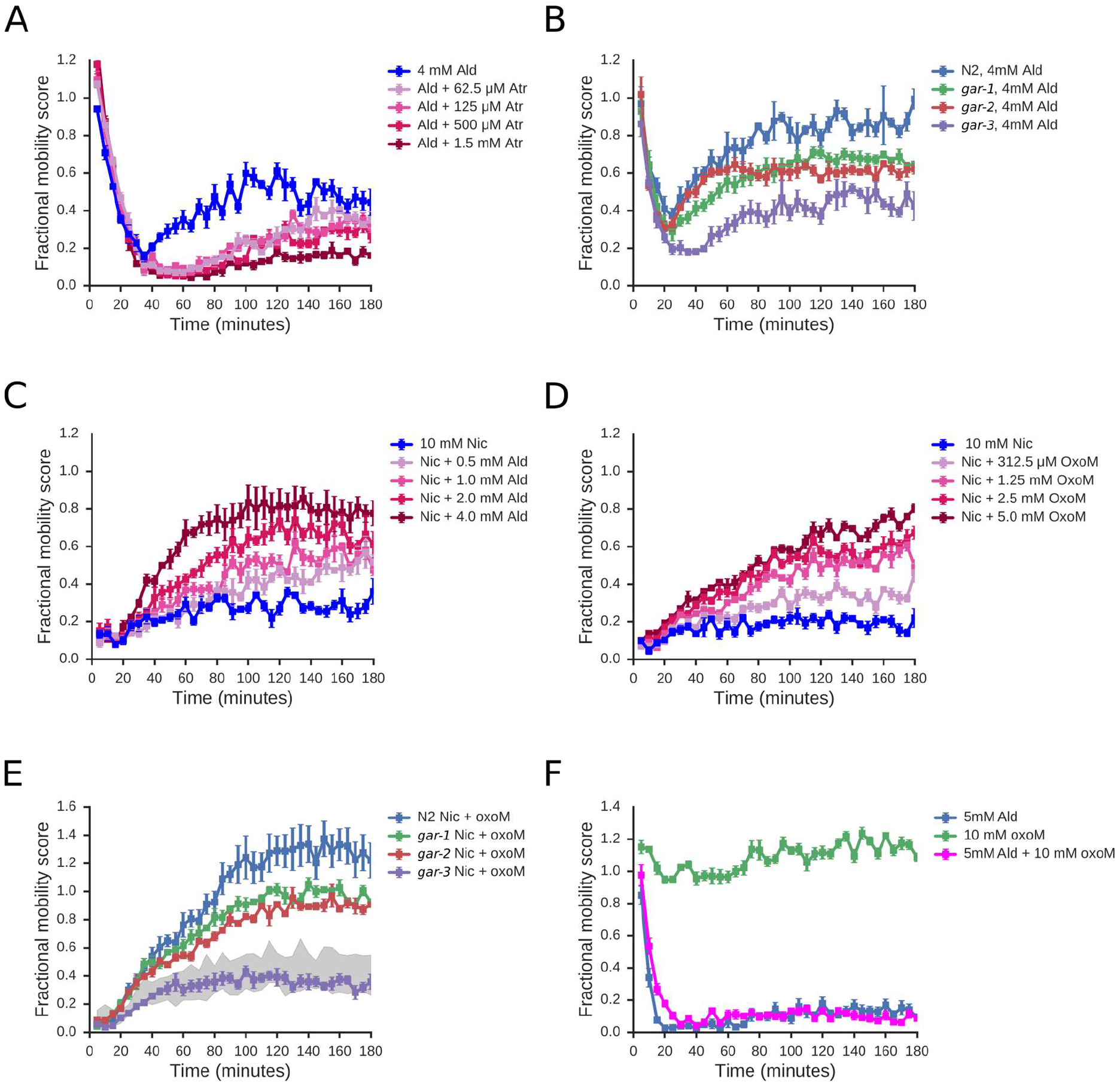
L1 cholinergic recovery can be modified by compounds with muscarinic activity. **A: The recovery from Ald-paralysis is suppressed by the mAChR antagonist Atr.** L1 worms were treated with 4mM Ald along with a range of doses of Atropine (Atr); FMS was measured every 5mins over a 3hr time course. **B: *gar-3* mutants show greatly reduced recovery from Ald-induced paralysis.** Wild-type animals, or worms homozygous for mutations in either *gar-1, gar-2*, or *gar-3* were exposed to 4mM Ald and FMS measured across a 3hr time course. **C: Aldicarb induces recovery from nicotine-induced paralysis in L1 larvae.** Various concentrations of aldicarb were combined with a concentration of nicotine sufficient to cause sustained paralysis (10 mM, shown in blue). Increasing concentrations of aldicarb stimulated increasing degrees of recovery from paralysis. **D: The mAChR-specific agonist OxoM similarly stimulates recovery from Nic-induced paralysis.** L1 animals were treated with 10mM Nic combined with a range of concentrations of OxoM. **E: OxoM-induced recovery requires *gar-3:*** Wild-type animals, or worms homozygous for mutations in either *gar-1, gar-2*, or *gar-3* were exposed to 10mM Nic and 10mM OxoM and FMS measured across a 3hr time course. The shaded portion represents the range of responses of N2, *gar-1, gar-2* and *gar-3* to nicotine, in the absence of OxoM. **(A-E)** All data are means of 4 independent experiments, normalized to no drug controls. Error bars show standard error of the mean. **F: OxoM-does not induce recovery from Ald paralysis in adult worms: requires *gar-3:*** Adult worms were exposed to 5mM Ald, with or without 10mM OxoM. The presence of OxoM does not induce recovery from Ald-mediated paralysis in adult worms. Data are means of three independent experiments; error bars show standard error of the mean.

We thus propose that Ald causes a complex acute response in L1 worms because the increased ACh levels have effects on two separate neurosignalling pathways. The first phase of the acute response is a rapid reduction in movement due to activation of nAChRs. The second phase is a slower recovery due to activation of mAChRs which somehow relieve the nAChR-driven paralysis. To further test this, we examined whether we can drive recovery from Nic-induced paralysis with agonists of mAChRs. Nic can only stimulate nAChRs and has no activity on mAChRs (30). Consistent with our model, we find that we can induce recovery from Nic-induced paralysis either by adding Ald (Fig 5C) or the mAChR agonists, Oxotremorine M (OxoM) or Arecoline (Are) (Fig 5D; supplementary S3). To confirm that the same mAChR requirement apply to OxoM-induced recovery as in Ald responses, we tested the ability of each *gar* mutant to recover from Nic-induced paralysis in the presence of OxoM (Fig 5E). We found that *gar-3* is indeed the primary mAChR driving OxoM-induced recovery. Our data suggest that mAChR signalling can drive recovery from paralysis induced by sustained nAChR activation. Finally, we examined whether changes in mAChR signalling might underlie the difference in the ability of L1 and adults to recover from Ald paralysis. We find that stimulation of mAChR signalling by OxoM in adults cannot override the paralysis response (Fig 5F) and thus, changes in the mAChR signalling machinery might indeed account for the different responses across development.

Our data thus suggest that a model in which there is some kind of physiological cross talk between two types of ACh receptor, nAChRs and mAChRs. This model is summarized in Fig 6, and suggests that i) sustained activation of nAChRs alone results in long-term paralysis, ii) sustained activation of mAChRs alone has little effect on movement, but iii) dual activation of both nAChRs and the mAChR GAR-3, such as with Ald alone, or Nic + OxoM results in an initial phase of nAChR-driven paralysis followed by a slower mAChR-driven recovery. We conclude that the complex acute response of L1 worms to Ald is highly specific to the drug used and is not a generic stress or xenobiotic response. To examine whether this is true for other drugs, we next looked at the effects of a very different drug, KCN, on worm movement.

**Figure 6:**
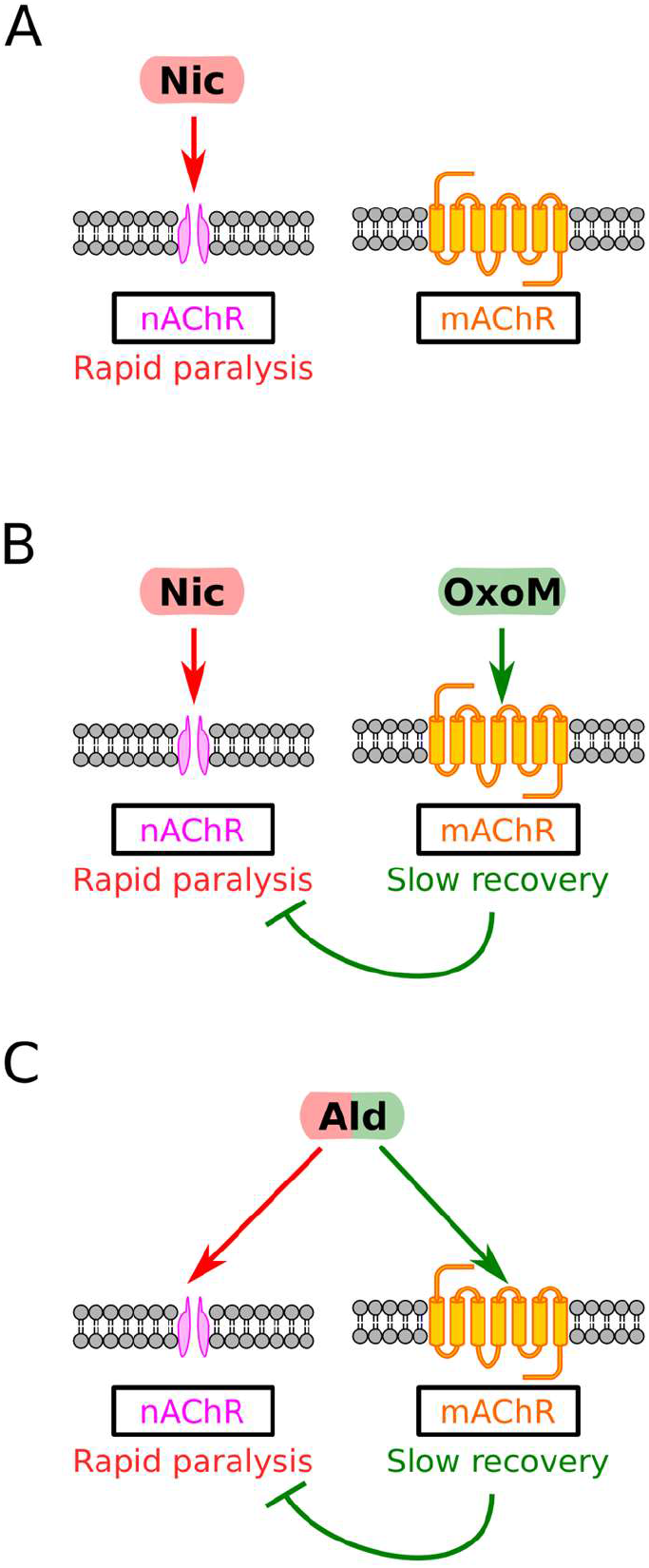
Conceptual model of cholinergic responses in L1 *C. elegans*. **A:** Nic alone causes paralysis through binding to nAChRs. **B:** The paralysis caused by the action of Nic at nAChRs in relieved by the action of OxoM at mAChRs. **C:** By elevating levels of ACh, Ald treatment leads to transient paralysis by activation of nAChRs, followed by recovery from paralysis, mediated by the action of ACh on mAChRs.

### *C.elegans* has a complex response to treatment with cyanide

Cyanide (KCN from here on) has a very well-characterized mode of action: it inhibits aerobic metabolism by blocking the mitochondrial electron transport chain (ETC) through binding to complex IV (reviewed in (31)). KCN is toxic to *C.elegans* following prolonged exposure — here we examined the acute responses of worms to KCN exposure.

We immediately noticed that the dose response at early time points were very unusual — while intermediate doses of KCN caused almost complete paralysis movement, high doses appeared to have little effect (Fig 7A). This was most pronounced in DRCs between 15-30minutes — by ~60 minutes the DRCs appeared much more normal, with increased effects being seen with increased dose.

**Figure 7.**
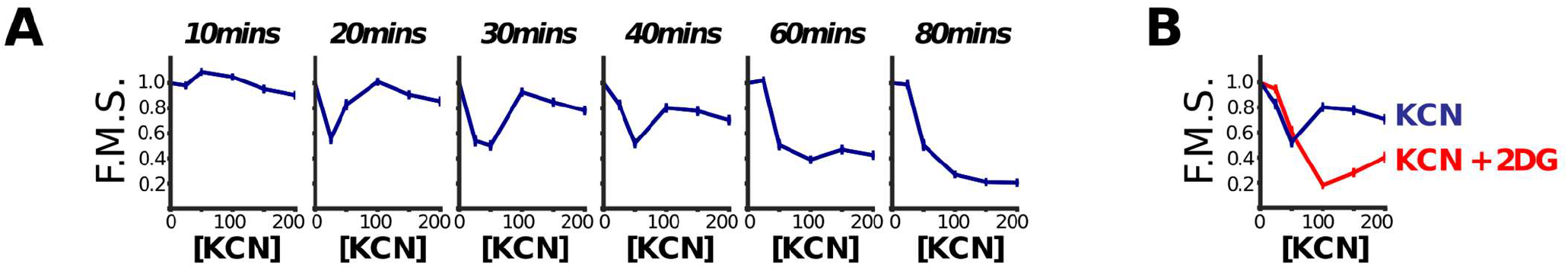
Dose sensitivity to KCN is complex. **A: L1 worms were treated with a range of KCN doses and their movement measured over time.** The Fractional Mobility Scores (FMS) are shown at various time points. At early time points the dose response curves are complex — intermediate doses cause more severe movement defects than high doses. **B: Decreased sensitivity to KCN at higher doses is dependent on a metabolic transition to glycolysis.** Inhibition of glycolysis with 2-dexoy-D-glucose causes higher-doses of KCN to be more toxic than lower doses, in contrast to the dose-response relationship of KCN alone. This suggests that the relatively mild effects of higher KCN concentrations seen in panel A is at least partially attributable to KCN stimulating a shift in metabolic pathways from oxidative phosphorylation to glycolysis. Data shown are at the 20 minute time point. **A-B:** Curves show mean of 3 repeats; error bars are standard error of the mean.

How can the effect of low concentrations of KCN be greater than the effect of high concentrations? We reasoned that this may be due to rewiring of metabolic pathways — worms might respond to high KCN concentrations by rapidly switching to use alternative metabolic pathways for generating ATP. An obvious candidate for such a pathway was anaerobic glycolysis which is the major pathway for ATP generation in anaerobic conditions in many animals — indeed we found that addition of 2-deoxy-D-glucose, an inhibitor of phosphoglucose isomerase, a critical enzyme in anaerobic glycolysis, caused a profound change in KCN DRCs. Worms can no longer continue moving in high doses of KCN when 2DG is also present — instead we find that increased concentrations of KCN result in decreased movement at all time points (Fig 7A). This suggests that the unusual DRCs seen for KCN alone were due to a shift to utilization of anaerobic glycolysis at early timepoints in high KCN (Fig 7B).

This underscores the importance of measuring drug responses across time — if we had only measured KCN DRCs at 80 minutes, they would have looked completely unremarkable. It also confirms that the way that worms respond to acute drug treatment is highly specific to the drug itself — while the aldicarb response involves crosstalk between nAChRs and mAChRs, the KCN response involves metabolic switches. The rapid responses of worms to compounds thus do not appear to be generic stress or xenobiotic responses but are specific responses to the specific drug effects. They reveal how the organism can rewire and adapt to sudden inhibition of a key pathway.

### Natural variation in acute drug response is very complex

Drug responses are known to vary within natural populations. In *C.elegans*, natural variation in drug responses has been seen for several drugs including abamectin and etoposide (32,33). In a previous study we compared the effects of KCN on two natural isolates, the N2 and CB4856 isolate (34). These differ by around 1SNP/800bp (35), a similar degree of variation as that between two human genomes. In our previous study we simply examined one time point (90 minutes after drug treatment) and only looked at a single larval stage, the L1 stage — we wanted to expand this analysis to compare the effects of KCN on N2 and CB4856 across development. We decided to examine how a range of concentrations of KCN affect movement of all 4 larval stages of either N2 or CB4856 and in each case to examine how this response evolves over 3hr (Fig 8).

**Figure 8.**
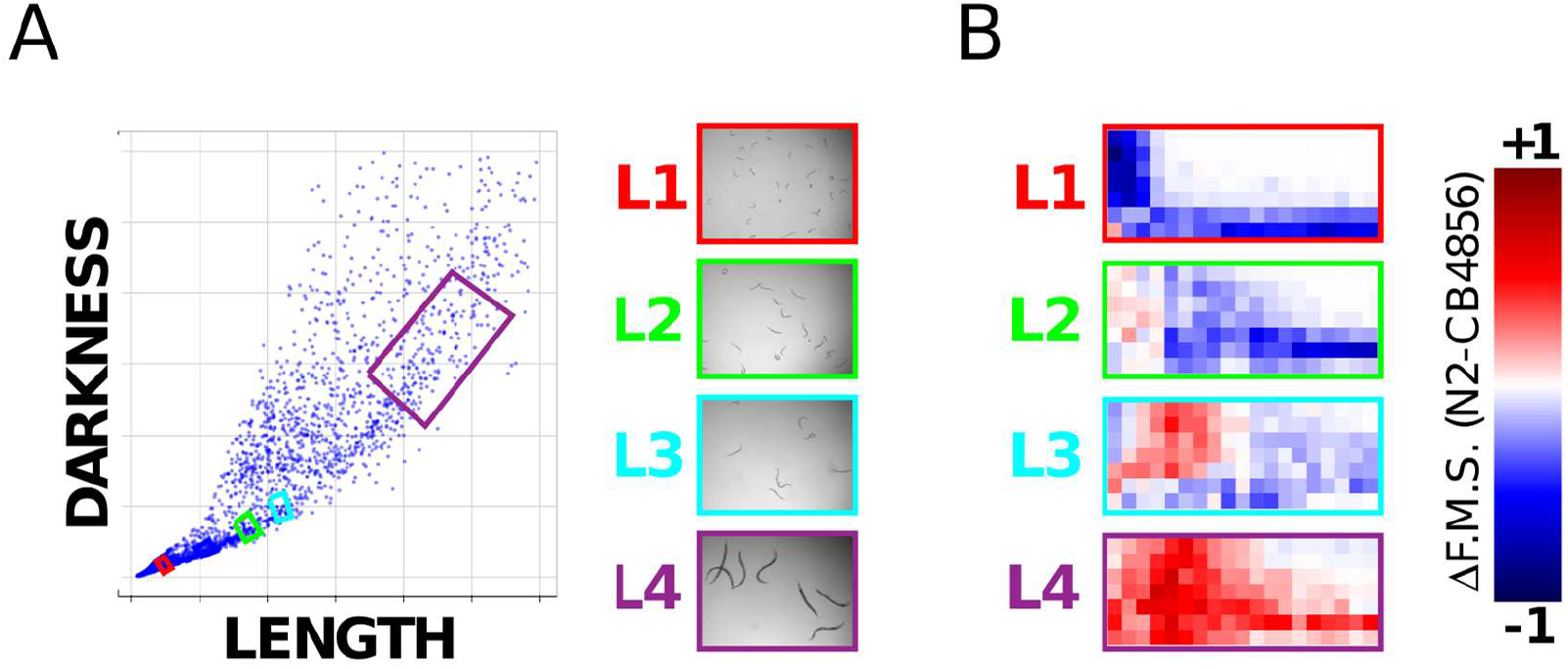
Variation in dose sensitivity to KCN changes across development. **A:** Worms of specific stages of either N2 or CB4856 isolates were purified using a worm sorter. Worms are sorted based on length and darkness — the exact windows chosen are indicated by coloured boxes and examples of the sorted worms shown. **B:** N2 or CB4856 worms of specific developmental stages were exposed to a range of doses of KCN and their movement measured over a 3hr time course. The fractional mobility score (F.M.S.) was calculated at each time point. The figure shows the difference in F.M.S. scores for N2 and CB4856 — blue colours indicate that N2 has a lower score, red colours that CB4856 is moving less.

The results are complex and striking. In keeping with our previous study (34), we find that N2 are more affected by KCN at the L1 stage. However, we find the exact reverse at the L4 stage — CB4856 are substantially more sensitive here. At the L3 stage, the picture is even more complex — while KCN appears to affect CB4856 more rapidly (CB4856 L3 animals are more severely affected at early timepoints), the same dose of KCN has a more severe effect on N2 later timepoints. Taken together, these data show clearly that comparisons of drug sensitivity in natural populations are complex. It is impossible to make a blanket statement that isolate A is ‘more sensitive’ than isolate B — assessing the effect at a different dose, a different timepoint or a different developmental stage could give precisely the reverse conclusion. These data underscore the need to measure drug responses as comprehensively as possible before making general conclusions — without measuring a range of doses and capturing these responses across time, we would have missed much of this complexity.

## Discussion

Drugs have extremely rapid effects on their targets — they can switch on or off their target within minutes even in a whole animal. What is the effect on the animal and how does it respond? We looked at the effects of drugs on *C.elegans* movement to measure acute responses to drugs in a whole animal and find several unexpected results that have important implications for drug screens in animals.

First, we found that around half of all the bioactive compounds that we found to affect *C.elegans* movement show a complex response: the worms are rapidly affected, but after an early phase of near-paralysis, the animals recover movement — this recovery may be transient or longer lived. In some cases, like the responses to compound shown in Fig 3E-F, we see multiple phases of paralysis and recovery over time — studying these complex acute responses of worms over time will drive exciting new insights. That worms can recover from the rapid effects of many drugs indicates that the worms are responding in some way. We did not expect to see this kind of recovery so frequently and to such a wide range of drugs and treatments, or responses of such complexity — that surprise drove this study.

Second, at least in the examples we examined, the mechanisms underlying complex acute responses appear to be drug specific and not due to a generic stress or xenobiotic response. For example, recovery from paralysis due to signaling by one set of acetylcholine receptors (AChRs) is due to signaling from a different set of AChRs; recovery from paralysis due to inhibition of aerobic respiration is driven by the use of anaerobic pathways. We see complex acute responses to a wide range of drugs and thus worms can respond to many different acute perturbations of their core networks by rewiring and modulating the activity of specific pathways. This suggests a tremendous complexity of homeostatic mechanisms and sensors and this molecular biology can be revealed through studying acute drug responses. For example, worms recover from paralysis induced by activation of nAChRs through signaling from mAChRs — this cross talk isn’t evident in chronic drug assays but can be easily measured using acute responses. Acute responses are thus excellent reporters for the rewiring of genetic networks in response to the effects of a drug.

Finally, we find that many factors affect acute drug responses. The acute response to the same drug can be very different at different developmental stages, at different doses of drug, or in different individuals — measuring only a single dose, a single time point, or a single developmental stage can greatly affect conclusions. For example, we found that L1 larvae recover readily from Aldicarb-induced paralysis but adults cannot recover. The recovery of L1s from Ald-induced paralysis is highly dose-dependent — L1s do not recover at 2mM Ald but recover readily at the higher concentration of 6mM (Fig 4A). This kind of complexity means one cannot simply say ‘*C.elegans* can recover from Ald paralysis’ or ‘*C.elegans* cannot recover from Ald paralysis’. We see a similar complexity when we look at natural variation in acute responses. We compared the sensitivity to KCN of two natural isolates of *C.elegans*, N2 and CB4856 and find a complex picture — while N2 is more sensitive at the L1 stage, CB4856 is more sensitive at the L4 stage. Again, we cannot make a blanket statement of ‘N2 is more sensitive than CB4856’— dose, time point, and developmental stage drive major differences in the drug responses and these factors are often neglected. For example a GWAS to identify QTLs that affect ‘drug sensitivity’ would likely find different variants depending on drug dose and developmental stage assayed. Crucially, this means that to really understand acute responses, we need to measure them over a range of doses and for each developmental stage. The simple assay we report here allows us to gather this kind of rich data.

Taken together, our results show that worms have complex acute responses to a wide variety of drugs and that these complex responses are due to the specific rewiring of the pathways underlying worm biology. Acute responses are sensitive readouts of the pathways that are required after rewiring — some of these are largely unexplored. For example, activation of mAChR signaling alone only has very subtle effects on worm biology (36–40) but acute responses reveal a clear effect of mAChR signaling in driving recovery from nAChR-induced paralysis. Genetic screens to find mutants with altered acute drug responses can thus dissect otherwise intractable pathways and acute responses can also be the basis for sensitive drug screens. Understanding the genetics that underpin complex acute responses like those shown in Fig 3 will reveal novel insights into how genetic networks rewire and change following insult.

Importantly, we also show that acute drug responses vary in many ways — different drug doses, different developmental stages, and different genetic backgrounds can give very different acute responses. Drawing broad conclusions about drug responses such as *‘C.elegans* recovers from Aldicarb-induced paralysis’ or ‘the N2 isolate is more sensitive to KCN than the CB4856’ is thus very difficult — different doses or different developmental stages often give completely different results. Studies on drugs in whole animals need to be aware of this complexity and interpret results with care. We believe that high throughput quantitative assays for drug responses of the kind we describe here are key — they allow researchers to build up a rich view of a drug response rather than focus on an arbitrary dose, time point or developmental stage. Studying the full complexity of acute responses opens up a whole new set of whole animal phenotypes for drug and genetic screens and will reveal how animals rapidly rewire their genetic networks in response to drugs, toxins, and environmental stresses.

## Materials and Methods

### Worm Maintenance and Strains

All worm stocks were maintained at 20°C on NGM agar plates seeded with *E.coli* strain OP50 as described elsewhere (41). In addition to the classical laboratory strain N2, we describe work using strains *unc-38* (*e264*), *lev-8* (*x15*), *lev-1* (*e211*), *unc-63* (*x26*), *unc-29* (*e1072*), *acr-8* (*ok1240*), *acr-16* (*ok789*), *gar-1* (*ok755*), *gar-2* (*ok520*) and *gar-3* (*gk305*). All strains were acquired from the Caenorhabditis Genetics Center.

To prepare L1 worms, animals were washed from the agar plates with M9 buffer (41) and rinsed in one buffer change of M9. L1s were then isolated by filtration though 11 μm nylon mesh filter plates (Milipore Multiscreen). L1 larvae for immediate use in drug response assays were diluted to approximately 1.2 worms per microlitre in either M9 buffer or modified NGM buffer (50 mM NaCl, 1 mM CaCl_2_, 1mM MgSO_4_, 200 mM KH2PO4, 50 mM K_2_HPO_4_).

To assay drug response over different developmental stages, worm populations were first developmentally synchronized by filtration of L1s, as described above. Filtered L1s were returned to fresh NGM agar 0P50 plates and incubated at 20°C until they had reached the required developmental stage (L1: no further incubation, L2: 18 hr, L3: 32 hr, L4: 42 hr, young adult: 54 hrs). Worms were then washed from the plates and rinsed as above. To further synchronize developmental stages, worms were dispensed into assay wells using a COPAS Biosort machine (Union Biometrica), which can selectively dispense individual worms on the developmentally-correlated criteria of size and optical density (L1 worms: 100 / well, L2: 80 / well, L3:, 50 / well, L4: 30 / well, young adult: 30 / well). Sorting parameters for each stage were determined empirically, as per the manufacturer’s instructions. The volume of liquid dispensed into the wells was determined and adjusted to 100 μl with buffer.

### Drug preparation

Solutions of potassium cyanide (Sigma Aldrich 60178-25G), sodium chloride (BioShop SOD002.205), cycloheximide (Sigma Aldrich C7698-1G) abamectin (Sigma Aldrich 31732) and 2-dexoxy-D-glucose (Sigma Aldrich D8375) were prepared fresh to twice the working concentration in M9 buffer, 1.6% DMSO. Stock solutions of aldicarb (Sigma Aldrich, 33386) were prepared to 1 M in DMSO. Solutions of levamisole hydrochloride (Acros Organics, 16595-80-5), oxotremorine M (Sigma Aldrich, O100) and mecamylamine hydrochloride (Sigma Aldrich, M9020) were prepared to 100 mM in water. These solutions were divided into single use aliquots and frozen. Stocks of arecoline hydrobromide (Acros Organics, 250130050) and atropine sulphate monohydrate (Sigma Aldrich, A0257) were prepared fresh on the day of use in M9 buffer. (-)-Nicotine (Sigma Aldrich, N3876) was stored frozen, undiluted in single use aliquots.

### Preparation of drug response assays

Assays were assembled in a final volume of 200 μl in flat-bottomed, polystyrene 96-well plates (Corning 3997) by combining 100 μl of worms in buffer with an equal volume of drug solution. To control for any confounding effects of drug solvent, all experiments were prepared to contain DMSO at 0.8 % v/v, regardless of whether the drug solution was prepared in DMSO or water. All assays were assembled in M9 buffer apart from the screen for compounds that modify the response to aldicarb, which were assembled in modified NGM, which we found to cause fewer problems of drug precipitation. After assembling the assay, plates were sealed with transparent, self-adhesive films before imaging. The point at which worms and drugs were combined marked the start of the assay.

### Preparation of RNAi assays

RNAi-by-feeding was performed as previously described (42). After 4 days, RNAi-treated and mock-treated worms were filtered to purify L1 worms and washed in two buffer changes of M9. Purified L1 samples were diluted to approximately 1.2 worms per microlitre and 100 μl samples were transferred to microtitre plates for use in mobility assays.

### Image acquisition, analysis and calculation of fractional mobility score

Images were acquired using a Nikon Ti Eclipse inverted microscope with 2x objective lens and DS-Qi1Mc camera. Image capture was automated in Nikon NIS-Elements AR software to capture two images of each well, separated by a 500 ms interval. This procedure was run in a 5 minute loop for the 3 hour duration of the experiments. A sample NIS-elements script is available as supplementary information.

Since our microscope control software stores files in a proprietary image stack format (Nikon ND2), we began our image analysis with a file format conversion process to convert nd2 files to a collection of 8-bit TIFF images, with structured filenames identifying the timepoint and well coordinates of each image. The first step of this process, defined in the Python file *export_nd2.py*, converted the ND2 file to a series of TIFF files, using the Open Microscopy Environment Bio-Formats tool (43). This conversion yields a series of 12-bit TIFF files with names encoding the timepoint and order of acquisition of each image. The second step of the conversion process invoked the ImageMagick package to convert these files from 12-bit to 8-bit TIFF, modify filenames to include the well location, rather than order of acquisition and group files for each timepoint into separate subfolders. This file structure forms the input for the image analysis script.

Our image analysis pipeline was prepared with the python programming language using the scikit-image library (44). All images were first processed with a Gaussian blur to reduce noise levels. Further image analysis consisted of two sets of processes to find moving and non-background, non-moving parts of the images.

To identify non-background, non-moving pixels, images were processed using a Sobel filter, which effectively produces the first derivative of the image, thus edges are highlighted and sharper edges appear brighter than shallow edges. An adaptive threshold was applied to the Sobel images to remove the background and yield a binary image. These binary images contained the outlines of the well and any objects, such as worms, within the well. Binary images were further processed with two iterations of binary closing to fill the pixels between sufficiently close edges.

This process was hand-tuned to fill in the area occupied by worms but minimize overfilling into the areas between worms. A circular binary mask was applied to remove the edges of the wells from each binary image and one of these images was designated as a reference image for each well (Figure 1A, Image B).

Pixels associated with movement were identified by calculating the absolute difference of the two consecutive images of a well (Figure 1A, Image A). A threshold was applied to the difference image to create a binary image (Figure 1A, Image B). This binary image contained the moving pixels from both parent images, showing double the number of movement-associated pixels. To correct for this, we multiplied the binary difference image by the reference image described above revealing only the non-background, movement associated pixels from a single frame (Figure 1A, Image C). Finally to identify pixels that did not move between wells, the reference image was multiplied by the complement of Image C (Figure 1A, Image D).

A fractional mobility score was calculated as the ratio of movement-associated pixels to total non-background pixels. This process is contained in the Python script *quantify_movement.py*. A second Python script, *quantify_movement_adult.py*, was used to analyse images of adult worms. This script contains an additional step to mask out objects below a certain number of pixels in size, to remove eggs from the analysis, since some chemicals stimulate egg-laying.

All our python scripts and an example dataset are available in our repository, https://github.com/fraser-lab-UofT/acute_assay

### Data analysis and curve-fitting

All drug response data shown were normalised by dividing by the fractional mobility score of drug-free control wells for each time point. Fitted lines were determined by least squares minimisation of either a three-parameter logistic function:

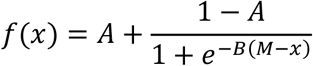

or an exponential function:

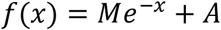

Where *x* denotes drug concentration and *M, A* and *B* denote parameters to be optimized. The choice of function was determined by the range of the dataset.

Proportional recovery scores were calculated using the formula:

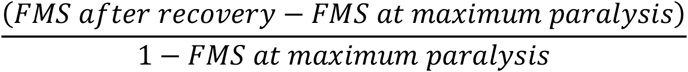

The value of FMS after recovery was calculated as the mean FMS over the period between 150 and 180 minutes. Maximum paralysis was determined as the minimum FMS value over the three hour course of the experiment.

---

**Figure S1:**
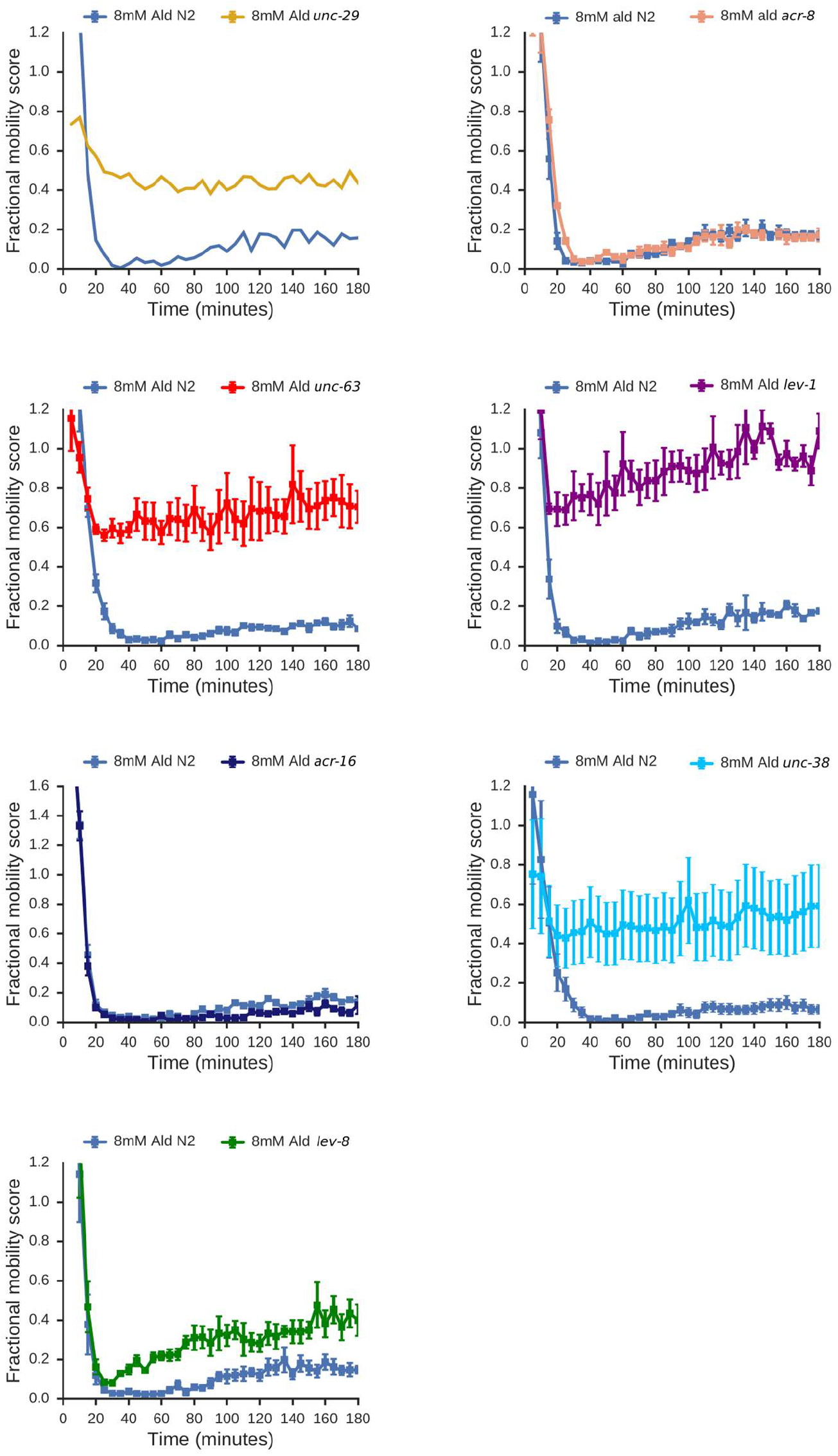
Response of nAChR subunit mutants to aldicarb in adult worms. Several nAChR subunits are required for wild-type sensitivity of L1 worms to 8mM aldicarb, including UNC-29, UNC-63, LEV1, LEV-8 and UNC-38. ACR-16 and ACR-8 are not required for Ald-sensitivity in adults. All data are means of 3 independent replicates. Error bars are standard error of the mean.

**Figure S2:**
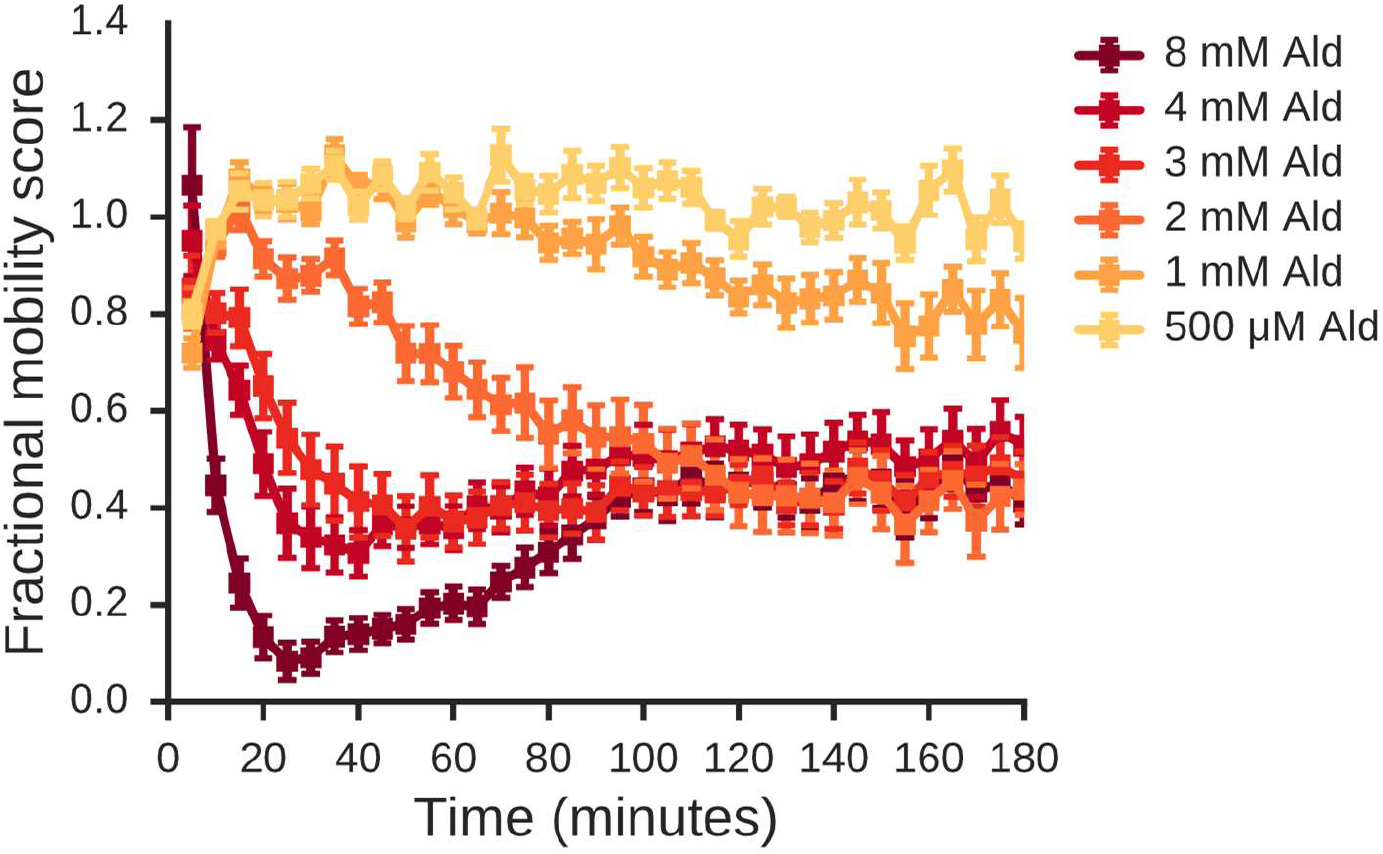
Further characterisation of mAChR recovery. Arecoline (Are), a mAChR agonist induces recovery from paralysis under 10 mM Nic in the same manner as OxoM. Data are means of 4 wells. Error bars are standard error of the mean.

**Figure S3:**
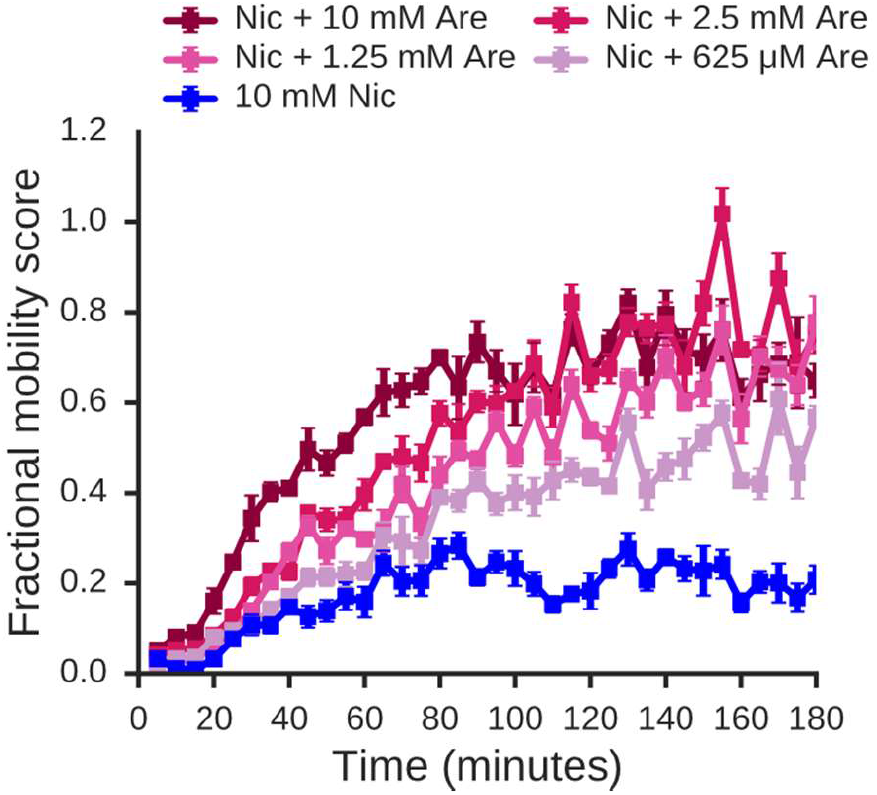
Time course of response of L1 worms to a range of Ald concentrations. Data are means of 4 independent experiments. Error bars are standard error of the mean.

